# Integrated Cross-Organ Transcriptomic Analysis Uncovers Conserved Gene Signatures Predictive of Allograft Rejection

**DOI:** 10.1101/2025.11.07.687316

**Authors:** H. C. Poorvi, P.K. Vinod

## Abstract

Long-term transplant success is limited by allograft rejection, a complex process traditionally studied on an organ-specific basis. To establish a unified framework beyond organ-specific studies, we performed a network-based systems biology analysis of transcriptomic data from 672 liver, kidney, and heart transplant biopsies to identify a conserved, pan-organ molecular framework of rejection. By constructing and comparing organ-specific gene co-expression networks, we identified a consensus, six-module immune cascade that captures the hierarchical nature of the alloimmune response. In addition, we also uncovered a highly conserved 24-gene cell cycle signature consistently upregulated in rejecting allografts, implicating cellular proliferation as a core feature of rejection pathology. From this framework, we derived a 172-gene immune signature and applied machine learning models to assess its predictive performance, achieving accuracy comparable to established benchmarks. We further refined this to a minimal, high-performance 20-gene immune signature (AUC > 0.96). Both the immune and cell cycle signatures demonstrated robust, pan-organ utility when independently validated in a lung transplant cohort (n=243). Collectively, these findings define a pan-organ molecular framework for rejection and highlight cell cycle dysregulation as a conserved hallmark, offering a foundation for standardized, cross-organ diagnostic platforms to improve allograft surveillance and patient outcomes.

## Introduction

Solid organ transplantation remains the gold standard therapeutic intervention for patients with end-stage organ failure, providing significant survival advantages and enhanced quality of life. However, long-term allograft outcomes remain suboptimal and variable across transplant types[1], with median survival rates of 14 years for kidney transplants, 14 for liver, 11 for heart and 5 for lung transplants[2]. This heterogeneity reflects the complex pathophysiology of allograft dysfunction, which encompasses ischemia-reperfusion injury, adaptive immune activation resulting in cellular and humoral rejection, and wound healing responses that replace functional parenchyma with fibrotic tissue[3].

The molecular characterization of transplant pathology has undergone rapid transformation through high-throughput genomic technologies and computational biology approaches, generating comprehensive datasets that hold promise for advancing precision transplant medicine. Multi-omics platforms enable systematic profiling of tissue microenvironments, providing detailed insights into host-recipient immune interactions and subsequent alloimmune cascades[3, 4]. Nevertheless, despite substantial genomic discoveries and biomarker identification efforts, clinical translation remains limited, with most molecular insights confined to research settings rather than routine diagnostic workflows[5].

Current molecular diagnostic approaches exhibit organ-specificity and lack standardization across transplant types, driven by assumptions that different organs demonstrate inherent biological heterogeneity in injury responses, immune recognition patterns, and repair mechanisms. Consequently, rejection monitoring relies on organ-specific gene panels that require independent validation studies and separate clinical implementation protocols for each transplant type, creating a complex and dissociated system. This limits diagnostic scalability, increases healthcare costs, and hinders systematic rejection monitoring across the transplant field[6].

However, accumulating evidence increasingly challenges this organ-specific framework. Studies have demonstrated cross-organ predictive utility of rejection biomarkers, with notable examples including the Molecular Microscope Diagnostics (MMDx) platform [7] and comprehensive pan-organ transcriptomic analyses[8], supporting the existence of shared biomarker capabilities across transplant types. These observations indicate that allograft rejection may be driven by conserved molecular mechanisms across different organs. The B-HOT (Banff of Human Organ Transplant)[6] gene panel, compiled through a comprehensive literature review of over 2000+ transplantation papers, represents a curated 770-gene collection encompassing all reported biomarkers across transplant studies. However, this gene panel compilation lacks systematic derivation and validation across organ types.

A systems-level transcriptomic analysis across multiple transplant types is still largely lacking. Such an approach could uncover shared pathophysiological networks and facilitate the discovery of universal rejection signatures, advancing both mechanistic understanding and standardized diagnostic development. Network-based systems biology methods offer a powerful methodological framework for addressing this knowledge gap by integrating high-dimensional omics data and deciphering complex pathophysiological processes underlying disease progression[9–11]. These approaches, encompassing gene co-expression networks, protein interaction networks, and regulatory networks, excel at identifying functionally coherent gene modules and their constituent elements. Through systematic analysis of these functional modules and their roles in disease pathogenesis, network-based methods enable mechanistic insights into the biological processes driving complex phenotypes such as transplant rejection.

In this study, we employed a systems biology approach to analyze transcriptomic profiles from liver, heart, and kidney transplant biopsies to identify conserved molecular mechanisms of pan-organ rejection **(Figure 1)**. Using network-based methods, we constructed organ-specific gene co-expression networks, identified rejection-associated modules within each organ, and performed cross-organ comparisons to derive consensus gene sets. We then evaluated the predictive utility of these consensus genes using machine learning models, refining it into a minimal 20-gene signature with robust performance across all three transplant types. This cross-organ framework not only captured the immune modules but also uncovered a conserved cell cycle regulatory program, implicating proliferative responses as a core and previously underappreciated feature of rejection pathology.

**Figure 1.**
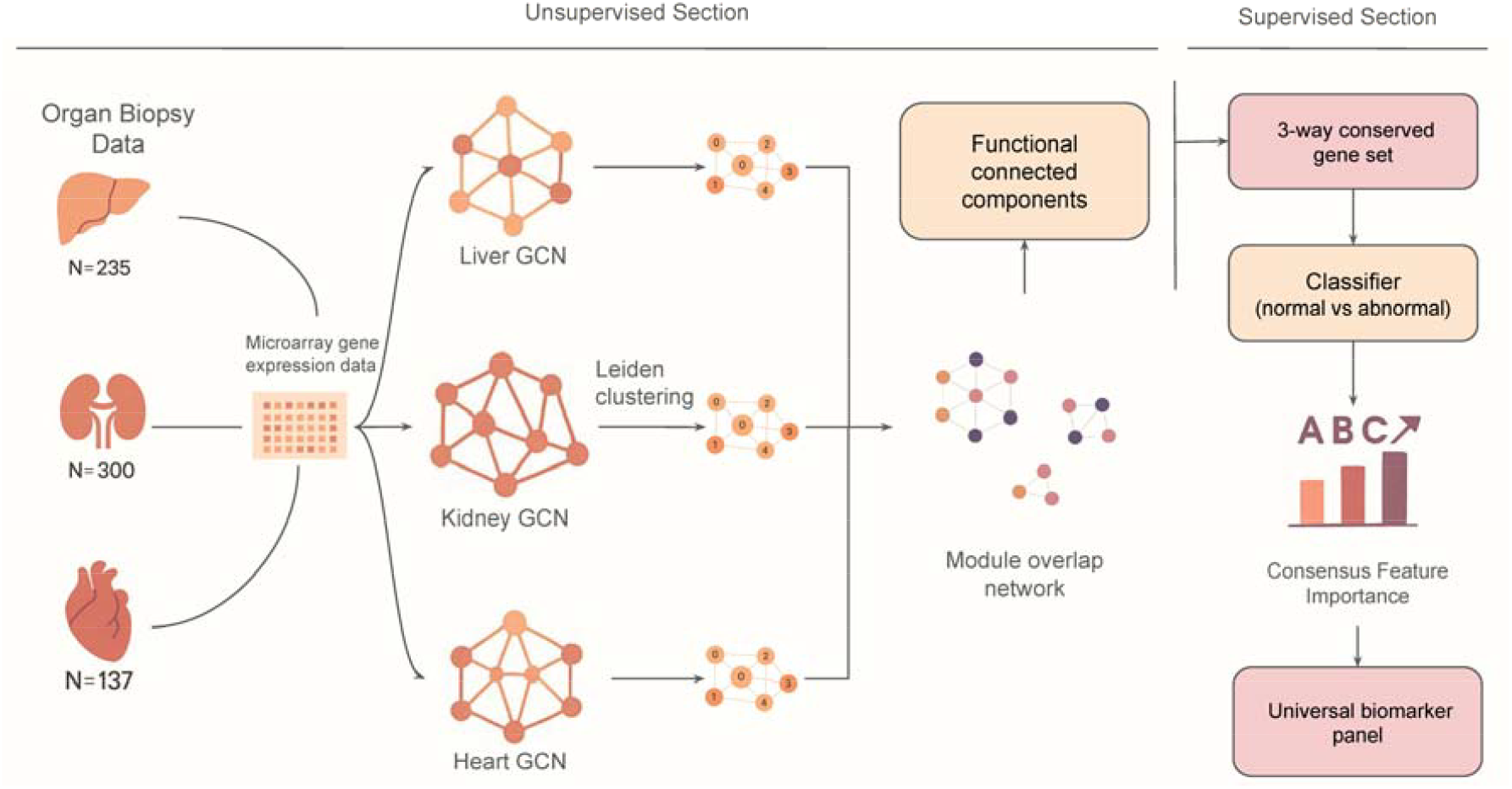
Overview of the Study Workflow: The study workflow consists of three main stages: (1) Unsupervised construction of co-expression networks (GCN) for liver, heart and kidney transplant cohorts; (2) Comparative network analysis to identify conserved functional modules across the three organs; (3) Development and validation of a pan-organ gene signature for allograft rejection using machine learning approaches.

## Methods

### Dataset Collection and Preprocessing

For our primary analysis, we utilized three publicly available microarray datasets from the Gene Expression Omnibus (GEO) database, representing liver (GSE145780), kidney (GSE192444), and heart (GSE272655) transplant biopsies[7, 12, 13]. These datasets encompassed a total of 672 samples across multiple rejection phenotypes and normal controls. To validate our findings, we employed an independent lung transplant dataset (GSE125004)[14], comprising 243 samples with diverse pathological states. All datasets had undergone standardized preprocessing procedures following established microarray analysis protocols. Raw expression data were processed using RMA [15], which included background correction, quantile normalization across arrays, and log2 transformation to ensure data comparability and reduce technical variability. The distribution of samples across different transplant organs and rejection phenotypes is summarized in **Table 1**.

**Table 1:** Dataset characteristics and sample distribution across transplant organs and rejection phenotypes used in this analysis. TCMR: T-cell mediated Rejection; ABMR: Antibody-mediated rejection; pABMR: Possible ABMR; pTCMR: Possible TCMR.

| Organ | GEO Accession | Phenotype | Sample Count | Total Samples |
| --- | --- | --- | --- | --- |
| Liver | GSE145780 | Normal | 129 | 235 |
|  |  | TCMR | 37 |  |
|  |  | Injury | 61 |  |
|  |  | Fibrosis | 8 |  |
| Kidney | GSE192444 | Normal | 175 | 300 |
|  |  | ABMR | 67 |  |
|  |  | TCMR | 21 |  |
|  |  | pABMR | 12 |  |
|  |  | pTCMR | 6 |  |
|  |  | Mixed Rejection | 19 |  |
| Heart | GSE272655 | Normal | 97 | 137 |
|  |  | ABMR | 12 |  |
|  |  | pABMR | 11 |  |
|  |  | TCMR | 10 |  |
|  |  | Mixed | 7 |  |
| Lung<br>(Validation dataset) | GSE125004 | Normal | 167 | 243 |
|  |  | Rejection | 24 |  |
|  |  | Late-inflammation Atrophy | 36 |  |
|  |  | Early Injury | 16 |  |

### Gene Co-expression Network Construction

Gene co-expression networks were constructed independently for each organ dataset to capture patterns of coordinated gene expression[16]. First, genes with mean expression intensities below a threshold of 3.0 were excluded to minimize technical noise. Next, variance-based filtering was applied to retain the top 25% most variable genes, focusing the analysis on genes that exhibit substantial expression changes across different sample conditions. Pairwise Spearman correlation coefficients were then calculated between all retained genes, with p-values adjusted using the Benjamini-Hochberg false discovery rate (FDR) correction.

To construct a high-confidence network suitable for downstream analysis, we applied a correlation threshold of |r| > 0.8. This cut-off was chosen to capture strong co-expression relationships that are more likely to reflect biological coordination. By taking this conservative approach, we minimized false positive interactions while retaining the robust gene-gene associations, which are expected to have mechanistic relevance in transplant rejection. The resulting interactions were represented as undirected weighted graphs, where nodes correspond to genes and edges reflect the strength of co-expression.

### Module Identification and Similarity Network

Leiden graph-based clustering[17] was applied to each co-expression network using a resolution parameter of 0.2 to identify functionally coherent gene modules. To assess the conservation of modules across organ types, a module similarity network was constructed, where each node represents a gene module identified within a specific organ. The similarity between any two modules was quantified using the Jaccard index, defined as the size of the intersection between two modules divided by the size of the union of the same two modules. A weighted edge with this value was drawn between them. The topology of this network was then analyzed to identify connected components, each delineating a “conservation group”. These groups comprise modules from two or more organs that are interconnected by significant gene overlap, representing the primary units of molecular conservation that capture both pairwise and three-way conserved regulatory programs.

The identified conservation groups were further validated through systematic pairwise and three-way comparisons. For pairwise analyses, a one-sided hypergeometric test was used to determine whether the gene overlap between two modules was greater than expected by chance. Fold enrichment was calculated as the ratio of observed to expected overlap. The expected overlap under random chance was computed as E[overlap] = (n□ × n□) / N, where n□ and n□ are module sizes and N is the number of genes in the co-expression network. For three-way comparisons, the analysis was framed as a pairwise statistical test, where one module was evaluated for significant overlap against the intersection of the other two modules. Functional enrichment analysis for conserved modules (identified through pairwise and three-way overlaps) was performed using Enrichr[18, 19], incorporating KEGG pathways, Reactome pathways, and Gene Ontology (GO) biological process databases.

### Performance benchmarking

Rejection-associated transcripts (RATs)[7, 20] represent a curated set of mRNAs whose expression levels are significantly altered during organ allograft rejection. Initially derived and extensively characterized in kidney transplant biopsies, RATs have since demonstrated generalizability and utility in the molecular assessment of rejection in other transplanted organs. RATs are categorized based on their association with specific rejection phenotypes: Universal RATs, TCMR-selective RATs and ABMR-Selective RATs. The Universal RATs exhibit the strongest statistical associations with rejection overall and comprise a comprehensive set of 417 transcripts, of which 350 were present in our datasets. Given their robust cross-organ applicability, RATs were used as the standard benchmark for evaluating the performance of the proposed biomarkers.

### Machine Learning Model for Rejection Prediction

We implemented a binary classification framework to distinguish normal transplant biopsies from pathological biopsies (i.e., those exhibiting rejection or injury) using a set of pan-organ conserved features. This approach directly addresses the primary clinical question while mitigating the class imbalances inherent in multi-class scenarios **(Table 1)**. Models were trained and tested on liver (129 normal vs 106 pathological biopsies), kidney (175 vs 125), and heart (97 vs 40) transplant cohorts and subsequently validated on an independent lung transplant dataset (167 vs 76).

To ensure robust and interpretable results, we employed three distinct machine learning algorithms. First, a Random Forest (RF) classifier with 100 estimators was selected for its strong predictive performance and built-in feature ranking capabilities. Second, a linear Support Vector Machine (SVM) classifier was used to test the linear separability between the classes. Finally, a Lasso-regularized logistic regression (lambda=0.1) was implemented to perform simultaneous classification and feature selection, thereby identifying a minimal set of highly predictive genes. Model performance was evaluated using a stratified 10-fold cross-validation scheme[21], with results reported as mean accuracy and AUC along with their corresponding standard deviations across folds.

### Feature Importance Analysis for Identifying Key Rejection Biomarkers

To identify the most discriminative genes for the pathological condition, we performed a post-hoc feature importance analysis. A consensus ranking of features was derived by integrating three complementary approaches. First, we extracted the built-in importance scores for each gene from the best-performing model. Second, we calculated permutation importance by measuring the decrease in AUC following random permutations of features. Third, SHAP (SHapley Additive exPlanations)[21] values were computed to quantify each feature’s contribution to model predictions. The scores from all three approaches were normalized to a [0,1] range and averaged to create the final consensus ranking, yielding a prioritized list of genes associated with the pathological condition.

## Results

### Network Topology and Modular Organization of Co-Expression Networks

The gene co-expression networks constructed for each transplant cohort revealed distinct topological architectures, reflecting the organ-specific regulatory landscapes. The networks varied in both size and complexity, with the kidney network exhibiting the greatest connectivity comprising 1,329 nodes and 26,638 edges followed by the heart (916 nodes, 11,642 edges) and liver (871 nodes, 6,282 edges) networks. Topological metrics further highlighted these differences: the kidney network demonstrated the highest clustering coefficient (liver: 0.464, kidney: 0.551, heart: 0.469), and a network density nearly twice that of liver (0.030 vs. 0.017) and substantially higher than heart (0.024).

Despite these global differences, all networks displayed highly modular architectures characteristic of biological systems. Each network comprised 43 to 55 connected components, indicating the presence of multiple distinct regulatory subnetworks operating within each organ. Across all three organs, the largest connected component consistently contained 607 to 670 nodes (69.4-77% of total nodes), likely representing core transcriptional programs common to transplant rejection responses. The remaining smaller components likely capture more cell-type-specific regulatory programs.

Leiden clustering revealed an inverse relationship between network connectivity and the number of identified modules. The highly connected kidney network was organized into 10 distinct modules, whereas the sparser liver and heart networks yielded 13 and 15 modules, respectively. This pattern suggests that increased network integration may lead to the formation of fewer but more cohesive functional modules, consistent with network theory predictions that highly connected biological networks tend to organize into larger, more integrated regulatory modules [22, 23].

### Cross Organ Module Conservation Patterns

A cross-organ comparative analysis of the co-expression networks revealed substantial overlap among the high-confidence genes across the three transplant types, suggesting the presence of shared transcriptional programs. A total of 648 genes were shared between at least two organs, with 280 genes found to be common across all three: liver, kidney and heart **(Figure 2A)**. This conserved gene set provided the basis for a more granular, module-level comparison to identify specific functional units preserved across organs.

**Figure 2:**
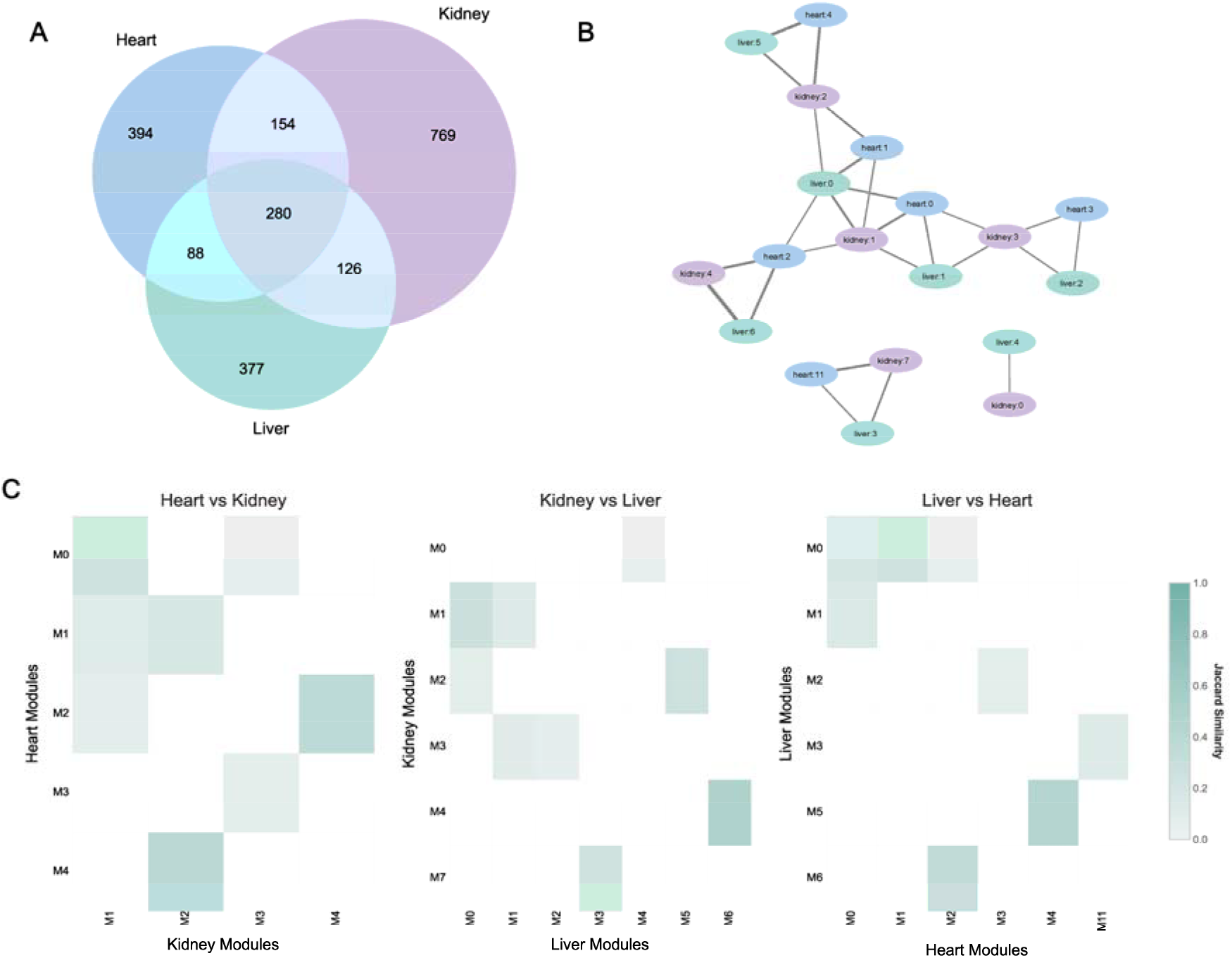
Cross-Organ Network and Module Conservation. **(A)** Venn diagram showing the overlap of network genes between liver, kidney, and heart. **(B)** Module similarity heatmaps showing pairwise overlaps between modules: Heart vs. Kidney, Kidney vs. Liver, and Liver vs. Heart, quantified using the Jaccard index. **(C)** Module similarity network, where nodes represent modules (colored by tissue type) and edges are weighted by their Jaccard indices, illustrating cross-organ module conservation.

To map the relationships between modules, we quantified similarity using the Jaccard index. Heatmaps of pairwise module comparisons showed distinct patterns of high similarity, where specific modules from one organ strongly mapped to one or more modules in another, indicating functional correspondence **(Figure 2C)**. To visualize this complex web of relationships, we constructed a module similarity network. **(Figure 2B)**.

The topology of the module similarity network revealed three distinct connected components, representing the primary units of molecular conservation across organs **(Figure 2B)**. These three conservation groups were designated as BC1, BC2 and BC3 **(Supplementary Table 1)**. The network topology also provided strong evidence for three-way conservation (triangular motifs), particularly within the major conservation group, BC1. Within BC1, we observed multiple triangular motifs, where modules from the liver, kidney, and heart were all interconnected. The presence of these highly interconnected triplets indicates that a shared set of genes is active in distinct modules across all three organs, reflecting a coordinated pan-organ regulatory program. BC1 comprised 14 modules from all three organs (*liver:0*,*1*,*5*,*6*,*2; kidney:1*,*2*,*3*,*4; heart: 0*,*1*,*2*,*3*,*4)* and contained a set of 452 conserved genes. BC2 also showed three-way conservation, forming a smaller, distinct group of three modules *(liver:3; kidney:7; heart:11)* with 24 shared genes. In contrast, BC3 represented a bi-organ conserved program, consisting of 2 modules *(liver:4; kidney:0)* shared only between the liver and kidney, with 23 conserved genes. Together, these conserved groups accounted for 499 unique genes exhibiting bi- or tri-organ conserved expression patterns, defining a common molecular response to transplantation.

### Functional characterization of the Conserved Programs

Functional analysis of the three conservation groups revealed distinct biological themes. BC1 represented the primary immune response, whereas BC2 and BC3 captured a conserved cell cycle program and a shared metabolic response, respectively. Within the BC1 similarity network (Figure 2B), we identified six core triangular motifs designated as functional subgroups (C1-C6) **(Supplementary Table 2)**, each comprising interconnected modules from liver, kidney and heart. Pathway enrichment analysis was performed for each subgroup **(Figure 3A)**, with fold enrichment scores calculated relative to random expectation and ranked accordingly **(Figure 3B)**.

**Figure 3:**
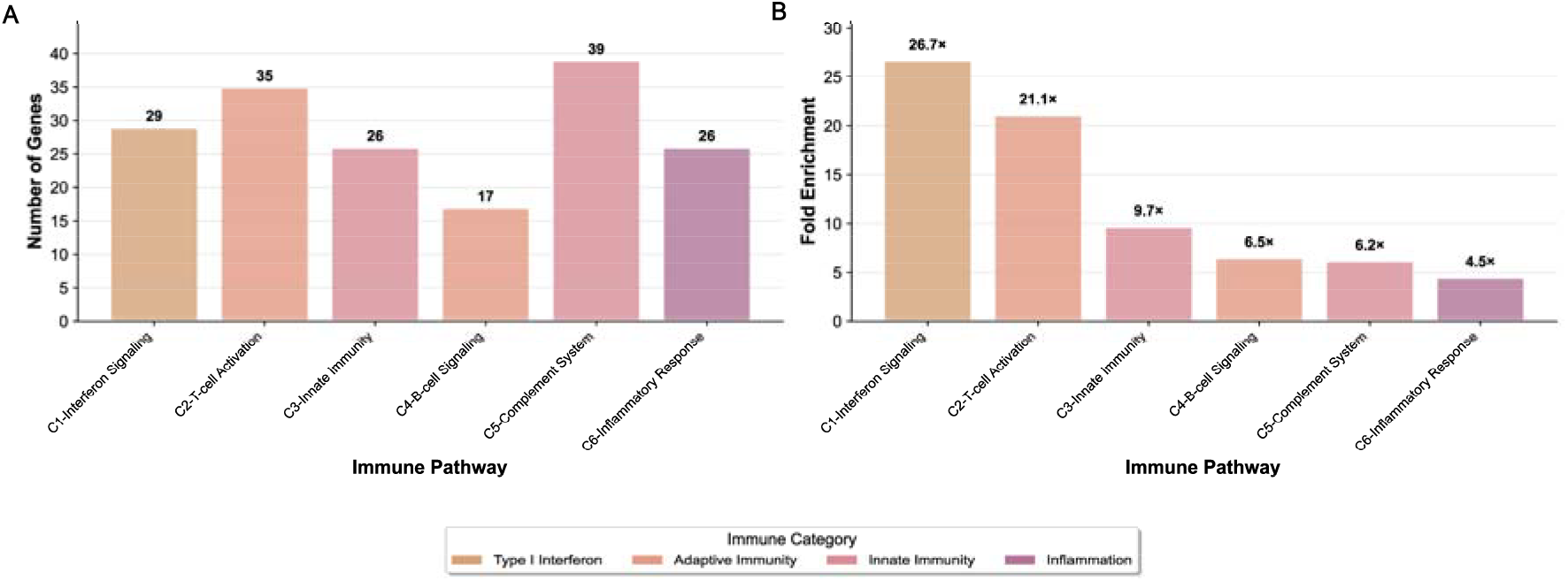
Three-way conserved immune pathway subgroups. **(A)** A bar graph showing the number of genes in each of the six three-way conserved subgroups (C1-C6), categorized by their functional enrichment. **(B)** A bar graph depicting the fold enrichment for each subgroup, highlighting the statistical significance of their conservation across liver, kidney and heart.

The C1 subgroup (Liver:6 + Kidney:4 + Heart:2) exhibited the highest fold enrichment (26.71) and comprised 29 genes central to interferon signalling. This subgroup included key regulatory components (STAT1, IRF1, and ISG15), chemokines (CXCL9/10/11), antiviral sensors (IFIH1, DDX60), and essential MHC class I processing components (HLA-B/E/F, TAP1, PSMB8/9). Functional enrichment analysis confirmed strong overrepresentation of Reactome pathways: Interferon Signaling and Interferon Alpha/Beta Signaling; KEGG pathways: Antigen Processing and Presentation, and GO term: Response to Type I Interferon, establishing C1 as a central interferon signaling module.

The C2 subgroup (Liver:5 + Kidney:2 + Heart:4) showed 21.1-fold enrichment and comprised 35 genes defining cytotoxic T lymphocyte machinery, including CD8A, PRF1 and GZMA, T-cell receptor signaling components (CD3D, LCK, ITK), co-stimulatory receptors (CD27, CD96), and transcriptional regulators (EOMES, RUNX3). These genes were significantly enriched in Reactome pathways: Adaptive Immune System and TCR Signaling; KEGG pathways: T Cell Receptor Signaling and Th1 and Th2 Differentiation, and GO term: T Cell Activation processes, collectively defining C2 as the T-cell cytotoxic effector module.

Moderately conserved subgroups C3-C6 exhibited distinct immune specializations. The C3 subgroup (Liver:0 + Kidney:1 + Heart:1), with 9.7-fold enrichment, captured innate immunity, comprising 26 genes including pattern recognition receptors TLR8 and CLEC7A and myeloid activation markers CD86 and CSF2RB. This subgroup was enriched for pathways related to Toll-like Receptor Cascades, Innate immune response, and Neutrophil extracellular trap formation pathways. The C4 subgroup (Liver:0 + Kidney:2 + Heart:1), with 6.5-fold enrichment, reflected B-cell signaling, including 17 genes such as CD38, PTPRC, and RAC2 and was enriched for B cell receptor signaling pathway, Lymphocyte activation, and Primary immunodeficiency pathways.

The C5 subgroup (Liver:0 + Kidney:1 + Heart:0), with 6.2-fold enrichment, captured 39 genes spanning classical complement components (C1QA/B/C), Fc receptors (FCGR3A), and macrophage markers (CD163), with strong enrichment for Complement and coagulation cascades and Phagosome pathways. Finally, the C6 subgroup (Liver:1 + Kidney:1 + Heart:0), with 4.5-fold enrichment, represented inflammatory resolution, featuring 26 genes such as anti-inflammatory mediator ANXA1, complement receptor C3AR1, and the immunoregulatory enzyme ENTPD1.

Together, these subgroups define a hierarchically organized immune surveillance network that reflects the sequential pathophysiology of transplant rejection. Initial ischemia-reperfusion injury activates innate immune pathways through the release of damage-associated molecular patterns (DAMPs) (C3), which subsequently trigger interferon signaling networks (C1) to coordinate immune activation. This response recruits and activates T-cells (C2), while engaging B-cell signaling (C4) and complement system activation (C5). Complement activation generates both tissue-bound and soluble inflammatory mediators, supported by macrophage-mediated responses. The inflammatory resolution pathways (C6) ultimately govern the clinical manifestation of the pathological condition.

Beyond the canonical immune conservation group, we identified a highly conserved 24-gene cell cycle program (BC2), consistently preserved across three organ modules (liver:3, kidney:7, heart:11). This subgroup contained key regulators of cell division, including the proliferation marker (MKI67), mitotic checkpoint proteins (BUB1B, TTK), regulatory cyclins (CCNA2, CCNB1), and DNA replication machinery (TOP2A, RRM2). Enrichment analysis confirmed significant overrepresentation of Reactome pathways: Cell Cycle, Mitotic and Mitotic Spindle Checkpoint, KEGG pathways: Cell Cycle, and GO term: Mitotic Sister Chromatid Segregation. Expression profiling demonstrated robust upregulation of the cell cycle genes in pathological samples across all organs **(Figure 4)**, highlighting cellular proliferation as a universal, previously underappreciated mechanism of rejection.

**Fig 4:**
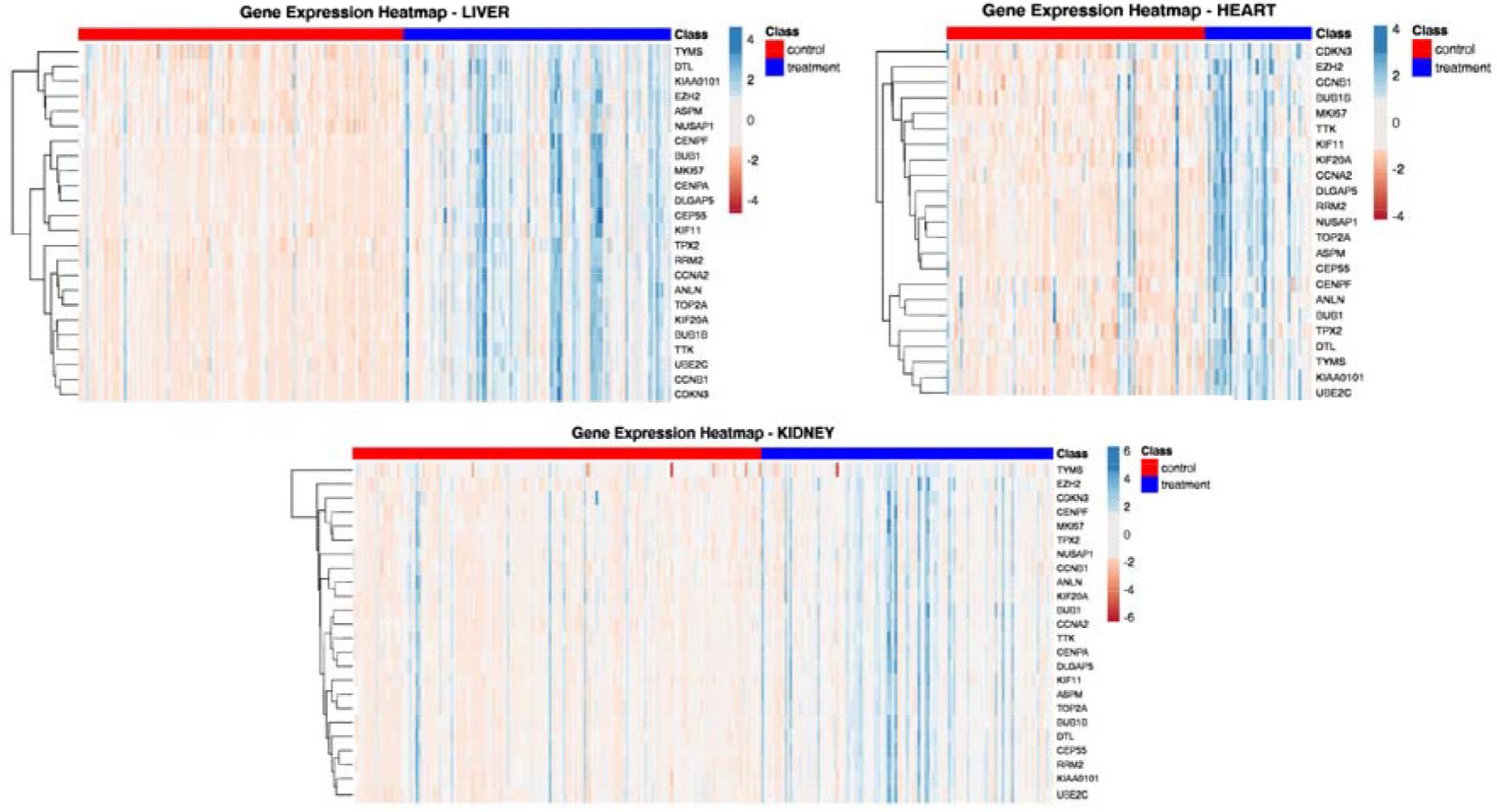
Gene Expression Heatmaps of Cell Cycle-Related Genes Across Organs. Heatmaps showing the expression levels of the 24-cell cycle-related genes (BC2) in heart, kidney, and liver biopsy samples.

A third conserved signature, designated BC3, comprised a 23-gene group shared between liver module 4 and kidney module 0. Pathway enrichment analysis implicated this group in core metabolic processes, showing significant overrepresentation of pathways related to amino acid and fatty acid metabolism. Enriched pathways included Metabolism of Amino Acids and Derivatives (Reactome) and Glycine, Serine, and Threonine Metabolism (KEGG). The conservation of this metabolic signature suggests a shared metabolic stress response characteristic of the allograft rejection process in these organs [24].

Together, these findings define a pan-organ molecular framework for transplant rejection that integrates canonical immune cascades with cell proliferation and metabolic dysregulation, offering novel insights into rejection biology and potential therapeutic targets.

### Predictive Performance of Gene Signatures Derived from Conserved Modules

We next evaluated the clinical utility of conservation groups (BCs) as diagnostic signatures for predicting pathological allograft behaviour. Supervised machine learning models were trained using the gene sets corresponding to the immune subgroups within BC1(C1-C6), and their ability to distinguish pathological from normal allograft biopsies was assessed across the three organ cohorts **(Table 2)**. Comparative analysis revealed that the interferon signaling subgroup (C1) and the T-cell cytotoxicity subgroup (C2) consistently achieved the highest classification performance. The C1 subgroup, which includes the established kidney rejection biomarker ISG15, performed well in heart (AUC = 0.973) and kidney (AUC = 0.952) datasets. In liver, the T-cell activation subgroup (C2) performed marginally better (AUC = 0.930) than C1 (AUC = 0.926). The macrophage-associated C5 subgroup also demonstrated strong cross-organ performance (AUC > 0.919), underscoring the role of M1/M2 macrophage balance as a universal determinant of rejection status[25]. Overall predictive performance was slightly lower in the liver cohort, likely due to the inclusion of samples with parenchymal injury in the pathological group, a factor absent in the more immune-specific rejection classifications for kidney and heart.

**Table 2:** Predictive performance of the six three-way conserved immune subgroups. The table reports the best classification performance achieved by each conserved immune signature (C1-C6) across three machine learning models: Random Forest (RF), Lasso Regression (Lasso), Support Vector Machine (SVM). Reported metrics include Accuracy (ACC) and Area Under the Curve (AUC), with corresponding standard deviations.

| Subgroups | Liver |  |  | Heart |  |  | Kidney |  |  |
| --- | --- | --- | --- | --- | --- | --- | --- | --- | --- |
|  | ACC | AUC | Model | ACC | AUC | Model | ACC | AUC | Model |
| C1 | 0.834<br>± 0.06 | 0.926<br>± 0.05 | RF | **0.906**<br>± **0.06** | **0.973**<br>± **0.04** | **RF** | **0.920**<br>± **0.06** | **0.952**<br>± **0.05** | **SVM** |
| C2 | **0.838**<br>± **0.03** | **0.931**<br>± **0.05** | **SVM** | 0.897<br>± 0.07 | 0.964<br>± 0.03 | RF | 0.913<br>± 0.05 | 0.950<br>± 0.04 | RF |
| C3 | 0.825<br>± 0.06 | 0.902<br>± 0.07 | RF | 0.883<br>± 0.09 | 0.939<br>± 0.07 | RF | 0.850<br>± 0.05 | 0.907<br>± 0.04 | RF |
| C4 | 0.788<br>± 0.05 | 0.848<br>± 0.06 | RF | 0.898<br>± 0.07 | 0.954<br>± 0.03 | SVM | 0.823<br>± 0.05 | 0.888<br>± 0.06 | RF |
| C5 | 0.851 | 0.919 | RF | 0.876 | 0.931 | Lasso | 0.887 | 0.938 | RF |
| | $\pm 0.08$ | $\pm 0.06$ | | $\pm 0.07$ | $\pm 0.06$ | | $\pm 0.06$ | $\pm 0.04$ | |
| C6 | 0.841<br>$\pm 0.07$ | 0.926<br>$\pm 0.05$ | RF | 0.877<br>$\pm 0.1$ | 0.95 ±<br>0.07 | RF | 0.837<br>$\pm 0.05$ | 0.886<br>$\pm 0.04$ | RF |

Although the individual immune subgroups demonstrated strong predictive power, each represents only one piece of a complex biological cascade. We hypothesized that integrating all the immune subgroups would yield better performance. Accordingly, we consolidated the six BC1 subgroups into a unified 172-gene immune signature. This pan-immune signature achieved performance comparable to the established 350-gene RATs panel, despite using less than half the number of features **(Table 3)**. Its consistent high-level performance across liver, kidney, and heart cohorts, despite heterogeneous sample distributions and varying rejection mechanisms supports the presence of shared molecular programs underlying pathological rejection.

**Table 3:** Benchmarking conserved gene signatures against the RATs standard. The predictive performance of different subsets of the conserved gene signature was compared with established Rejection Associated Transcripts (RATs) panel. Accuracy (ACC) and Area Under the Curve (AUC) are reported for Random Forest models evaluated independently on the liver, heart, and kidney transplant dataset.

| Gene Signature | Liver |  | Kidney |  | Heart |  |
| --- | --- | --- | --- | --- | --- | --- |
|  | ACC | AUC | ACC | AUC | ACC | AUC |
| RATs panel (350 genes) | 0.9190 ±<br>0.04 | 0.9914 ±<br>0.01 | 0.9167 ±<br>0.04 | 0.9696 ±<br>0.03 | 0.9417 ±<br>0.05 | 0.9712 ±<br>0.04 |
| 172-gene signature | 0.9150 ±<br>0.06 | 0.9803 ±<br>0.03 | 0.9100 ±<br>0.04 | 0.9473 ±<br>0.04 | 0.9269 ±<br>0.06 | 0.9694 ±<br>0.03 |
| 20-gene signature | 0.9324 ±<br>0.05 | 0.9893 ±<br>0.0 | 0.9100 ±<br>0.03 | 0.9552 ±<br>0.05 | 0.9565 ±<br>0.05 | 0.9659 ±<br>0.05 |
| 6-gene signature | 0.9232 ±<br>0.04 | 0.9542 ±<br>0.03 | 0.9000 ±<br>0.05 | 0.9583 ±<br>0.05 | 0.9132 ±<br>0.06 | 0.9663 ±<br>0.05 |

To identify the most informative genes within this 172-gene immune signature, feature importance was assessed using complementary approaches and a consensus ranking was derived across methods (see methods section). This analysis revealed a hierarchy of discriminative genes with KLRD1 emerging as the strongest predictor (consensus score = 0.684), demonstrating more than two-fold higher importance than the second-ranked gene FCGR1A (consensus score: 0.333) **(Figure 5A)**. Other top-ranked contributors included NKG7, CXCL11, CXCL9, and PRF1, followed by MSR1, EOMES, APOL3, and SAMD3. These genes were consistently upregulated across rejection samples from all three organs **(Figure 5B)**. When visualized collectively, the average expression pattern of the top genes (**Supplementary Table 3**) exhibited a clear separation between pathological and normal sample groups (**Figure 5C–E**), underscoring their strong diagnostic utility.

**Figure 5:**
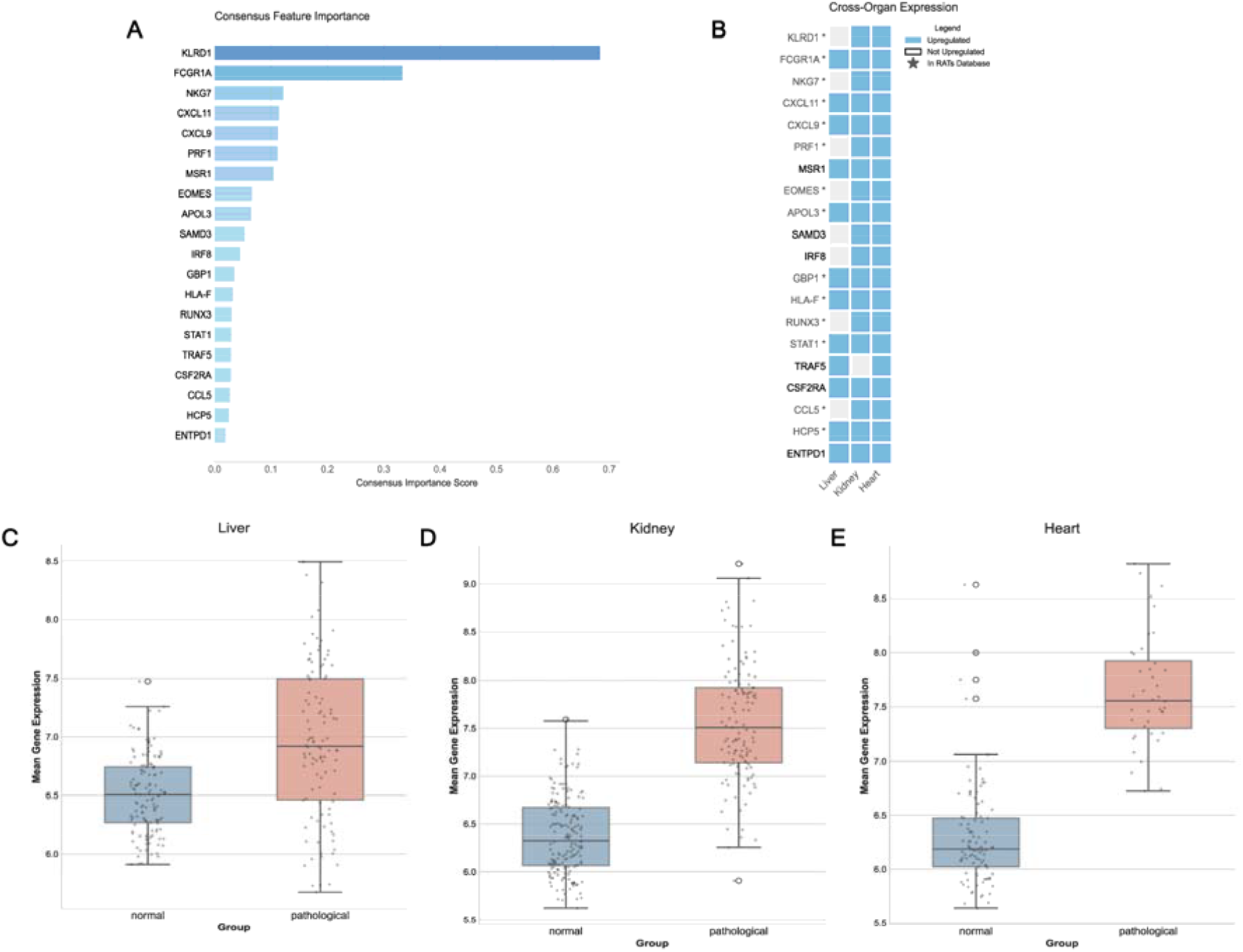
Identification and Validation of a Minimal Universal Rejection Signature. **(A)** Consensus feature importance scores for the top universal biomarkers. **(B)** The panel showing that each gene is significantly upregulated in rejecting samples across liver, kidney, and heart, along with their overlap with the established RATs panel (shown as *). **(C-E)** Boxplots showing the average expression levels of the top gene signature in liver **(C)**, kidney **(D)**, and heart **(E)** samples, illustrating clear separation between pathological and normal groups.

Systematic evaluation of model performance as a function of feature number showed that predictive accuracy plateaued at approximately 20 genes across all organ types **(Supplementary Figure 1)**. The top 20 consensus-ranked genes achieved nearly identical performance to the full 172-gene set (accuracy difference < 2%, AUC difference < 0.01), indicating that these features capture the essential discriminative information for rejection classification **(Figure 5C-E)**. Some of these genes also overlap with the RATs panel **(Figure 5B)**. While the 20-gene panel represents the point of peak performance, a minimalist 6-gene signature (KLRD1, FCGR1A, NKG7, CXCL11, CXCL9, PRF1) retained robust performance (AUC > 0.95 across all organs). This compact signature thus offers a practical, cost-efficient tool for rapid screening by capturing the core rejection signal with minimal features **(Table 3)**.

Further, we also evaluated the performance of the BC2 subgroup, comprising 24 cell cycle related genes, in distinguishing pathological from normal allograft biopsies. Predictive modeling revealed moderate yet consistent performance across all three organs (AUC = 0.84-0.85 in liver, 0.76-0.78 in kidney, 0.89-0.90 in heart). This independent predictive capability of this signature suggests that cell cycle dysregulation is a core biological process in rejection rather than a bystander effect of inflammation. The consistent upregulation of these genes in pathological samples across organs further support their direct involvement in the pathological process.

### Validation in an Independent Lung Transplant Cohort

To rigorously assess the pan-organ generalizability of these derived signatures, we performed external validation using an independent lung transplant biopsy dataset (GSE125004), an organ type not included in the discovery analysis. The 172-gene immune signature demonstrated high discriminatory performance (AUC = 0.994, accuracy = 0.959), while the 20-gene signature performed comparably well (AUC = 0.975). The minimal 6-gene set also maintained strong predictive capability (AUC = 0.939). The novel 24-gene cell cycle signature showed significant predictive performance (AUC = 0.746). The high predictive performance of these distinct biological signatures in a fourth solid organ provides compelling evidence that the molecular signals are not tissue specific. These findings suggest that our data-driven approach has uncovered a conserved, pan-organ molecular program underlying allograft rejection, supporting the development of a standardized diagnostic platform with broad applicability.

## Discussion

Solid organ transplantation is a transformative, life-saving therapy for end-stage organ failure, yet its long-term success is persistently challenged by allograft rejection. Historically, the pathophysiology of rejection has been investigated within an organ-specific paradigm. While this approach has yielded crucial insights, the shared molecular mechanisms driving rejection across different allografts remain less understood. This study addresses this critical gap by employing a systems biology framework to define a conserved molecular architecture of rejection across liver, kidney and heart transplants. By constructing and comparing gene co-expression networks, we demonstrate that despite distinct network topologies, common biological mechanisms orchestrate the alloimmune response, providing a foundation for a pan-organ diagnostic platform.

Previous pan-organ analyses have provided compelling evidence for a shared rejection program, most notably through the “Immunologic Constant of Rejection” (ICR) hypothesis [26]. This hypothesis posits that disparate forms of immune-mediated tissue destruction converge on a common set of final molecular pathways. This includes the activation of interferon-stimulated genes (ISGs), chemokine-mediated recruitment of cytotoxic cells, and the deployment of immune effector function genes like granzymes and perforin. Subsequent studies, like the “Common Rejection Module” (CRM) [27], and the comprehensive PROMAD atlas [8], corroborate the hypothesis through meta-analyses of differentially expressed genes. Our study adopts a systems-level approach to build a pan-organ framework for transplant rejection. By constructing organ-specific gene co-expression networks, we move beyond compiling shared gene lists to uncovering functionally conserved modules of co-regulated genes. Integrating these modules into a higher-order mechanistic framework reveals the hierarchical organization of the alloimmune response and highlights conserved biological processes.

The predictive power of this framework is demonstrated by the development of a minimal, high performance 20-gene signature that distils the complexity of rejection biology into a compact biomarker panel. Importantly, this signature was derived from network modules rather than solely from differential expression, representing conserved six immune subgroups(C1–C6) comprising 172 genes, capturing the core elements of alloimmune biology. These include transcriptional regulators (STAT1, IRF8, RUNX3, EOMES), effector molecules/regulators (PRF1, NKG7, KLRD1), interferon-inducible chemokines (CXCL9, CXCL11, CCL5), antigen-presentation markers (HLA-F, HCP5), signaling mediators (TRAF5, CSF2RA) and macrophage associated genes (FCGR1A, MSR1, ENTPD1), together delineating a coordinated immune circuit spanning activation, effector function, and resolution.

Within this framework, the C1-subgroup comprises IFN-γ-dependent transcription factor STAT1, whose inhibition reduces cellular rejection in mouse heart allografts[28]. It also activates the canonical JAK-STAT pathway and its downstream effector IRF1[26]. This leads to increased graft antigenicity through MHC upregulation (HLA-F, HCP5) and the production of CXCR3-ligand chemokines (CXCL9, CXCL11), canonical markers of effector lymphocyte recruitment across solid organs [26, 29]. The C2-subgroup reflects T-cell cytotoxic machinery (C2), comprising genes essential for cell-killing (PRF1) and markers of cytotoxic cells (NKG7 and KLRD1), which are established components of rejection signatures. The differentiation of these killer cells is directed by factors like RUNX3 and EOMES, whose roles in promoting effector function are well documented[30]. Concurrently, B-cell responses (C4) establish persistent alloimmunity through the generation of pathogenic donor-specific antibodies (DSAs), the hallmark of antibody-mediated rejection (AMR)[31, 32], which are further amplified by the complement and macrophage module (C5). Here, DSAs engage macrophages via the high affinity IgG receptor FCGR1A, creating a potent feed-forward loop of injury[33]. Finally, the inflammatory resolution pathways (C6), exemplified by ENTPD1, serve as a biological checkpoint determining whether the graft progresses to repair or pathological injury[34]. In addition to these validated biomarkers, the 20-gene signature also includes novel genes APOL3, SAMD3, TRAF5, and CSF2RA, highlighting potential targets for mechanistic studies and therapeutic exploration.

When benchmarked against the established 350-gene RAT panel[7, 20], our consolidated 172-gene pan-immune signature demonstrated comparable discriminatory power across liver, kidney, and heart biopsy datasets, despite comprising less than half the number of features. This cross-tissue performance supports the existence of a universal alloimmune program underlying rejection pathology. Further refinement to the 20-gene subset preserved high predictive accuracy (AUC > 0.96), underscoring that a small, mechanistically informed signature can robustly capture the central dynamics of graft injury.

Another notable finding is the identification of a highly conserved 24-gene cell cycle group as a universal signature of rejection. The consistent upregulation of these markers including marker of cell proliferation across organs with vastly different regenerative capacities highlights a previously underappreciated component of rejection. This proliferative signal likely represents the clonal expansion of infiltrating lymphocytes or a dysregulated, pathological repair response by the graft parenchymal and endothelial cells. In renal allograft rejection, a distinct population of proliferating Cd8+ T cells has been characterised by high expression of MKI67 and other proliferation markers[36]. The independent predictive capability of this signature suggests it is an important component of the pathological process, providing a direct molecular link between acute immune-mediated damage and the progression to rejection and fibrosis.

We acknowledge several limitations that provide avenues for future research. First, this analysis is based on retrospective data, underscoring the need for validation of the 20-gene signature in prospective cohorts. Second, our binary classification does not differentiate between specific rejection subtypes or non-rejection-related injury; future studies with larger, well-annotated cohorts are required to evaluate the signature’s discriminative power across these finer categories. A recent study in heart transplant biopsies identifies finer rejection archetypes and highlights the importance of distinguishing rejection from parenchymal injury[36]. While our current analysis is biopsy-based, this modular architecture lays the foundation for future extensions into extracting non-invasive biomarkers using blood or other biofluids and mapping to single-cell or spatially resolved transcriptomic data, enabling the dissection of cell-type specific contributions to rejection.

In conclusion, this study advances the understanding of allograft rejection by defining a universal molecular framework shared across liver, kidney, and heart transplants. Our approach captures the architecture of a conserved immune cascade and highlights cellular proliferation as a core component of rejection pathology. The distillation of these findings into a minimal, high-performance 20-gene signature represents a significant step toward a standardized, pan-organ diagnostic tool and provides a foundation for improved graft monitoring and targeted therapeutic strategies to improve long-term transplant outcomes.

## Acknowledgements

This work was supported by iHUB-Data, International Institute of Information Technology, Hyderabad, India. The funding body has no role in the study design and analysis.

## Competing interests

The authors declare that they have no competing interests.

## Notes

### Competing Interest Statement

The authors have declared no competing interest.

